# Oral drug repositioning candidates and synergistic remdesivir combinations for the prophylaxis and treatment of COVID-19

**DOI:** 10.1101/2020.06.16.153403

**Authors:** Malina A. Bakowski, Nathan Beutler, Emily Chen, Tu-Trinh H. Nguyen, Melanie G. Kirkpatrick, Mara Parren, Linlin Yang, James Ricketts, Anil K. Gupta, Mitchell V. Hull, Peter G. Schultz, Dennis R. Burton, Arnab K. Chatterjee, Case W. McNamara, Thomas F. Rogers

**Author notes:** These authors contributed equally to this work (Malina A. Bakowski, Nathan Beutler).

## Abstract

The ongoing pandemic caused by the novel severe acute respiratory syndrome coronavirus 2 (SARS-CoV-2), necessitates strategies to identify prophylactic and therapeutic drug candidates for rapid clinical deployment. A high-throughput, high-content imaging assay of human HeLa cells expressing the SARS-CoV-2 receptor ACE2 was used to screen ReFRAME, a best-in-class drug repurposing library. From nearly 12,000 compounds, we identified 66 compounds capable of selectively inhibiting SARS-CoV-2 replication in human cells. Twenty-four of these drugs show additive activity in combination with the RNA-dependent RNA polymerase inhibitor remdesivir and may afford increased in vivo efficacy. We also identified synergistic interaction of the nucleoside analog riboprine and a folate antagonist 10-deazaaminopterin with remdesivir. Overall, seven clinically approved drugs (halofantrine, amiodarone, nelfinavir, simeprevir, manidipine, ozanimod, osimertinib) and 19 compounds in other stages of development may have the potential to be repurposed as SARS-CoV-2 oral therapeutics based on their potency, pharmacokinetic and human safety profiles.

## Main Text

In early December of 2019 the severe acute respiratory syndrome coronavirus 2 (SARS-CoV-2) was identified as the cause of rapidly increasing numbers of severe pneumonia-like symptoms termed COVID-19^1^. Since then, SARS-CoV-2 has rightfully been given its pandemic status by the World Health Organization (WHO). As of May 4, SARS-CoV-2 has spread throughout the world causing a combined total of more than 3,407,000 confirmed infections and more than 238,000 reported deaths in 215 different countries^2^. Antiviral treatment options for COVID-19 are extremely limited. Many compounds, such as ribavirin, interferon, lopinavir-ritonavir, and corticosteroids that had been used in patients with SARS-CoV-1 and MERS-CoV, remain unvalidated for the newly emerged SARS-CoV-2^3^. Remdesivir, a nucleotide analog prodrug with broad antiviral activity that works as an RNA-dependent RNA polymerase inhibitor, has been reported to be effective against MERS-CoV and SARS-CoV-1 infections in animal models^4–6^. Repositioning of remdesivir as a treatment against SARS-CoV-2 infection has recently demonstrated positive clinical endpoints in a Phase 3 Adaptive COVID-19 Treatment Trial (median time to recovery shortened from 15 to 11 days)^7^ that justified emergency use authorization of remdesivir by the US Food & Drug Administration for treatment of hospitalized COVID-19 patients^8^. The effectiveness of the repurposed drug remdesivir, with the current formulation requiring administration by IV infusion, both highlights the importance of investigating pre-existing drugs to combat SARS-CoV-2 infections, and the need for discovery of new or supplemental therapies that result in greater clinical improvements and can be administered outside of a hospital setting (i.e. orally). The ReFRAME (Repurposing, Focused Rescue, and Accelerated Medchem) drug collection is an extensive drug repurposing library containing nearly 12,000 small-molecule drugs shown to be appropriate for direct use in humans^9^ and provides a rich resource to discover new treatments that may be used as additional monotherapies or even in combination with remdesivir to further enhance efficacy and reduce drug resistance potential.

### High-throughput cell-based phenotypic assay against SARS-CoV-2

To identify compounds that could inhibit entry or replication of SARS-CoV-2 in human cells, we developed a high-content imaging (HCI) 384-well format assay using HeLa cells expressing the human SARS-CoV-2 receptor, the angiotensin-converting enzyme 2, or ACE2 (HeLa-ACE2). In this assay HeLa-ACE2 cells are infected with SARS-CoV-2 virus in the presence of compounds of interest, and viral infection is quantified 24 hours later (Fig. 1A). The assay relies on immunofluorescent detection of SARS-CoV-2 proteins with sera purified from patients exposed to the virus, which together with host cell nuclear staining allows for quantification of the percent infected cells in each well (Fig. 1B).

**Fig. 1.**
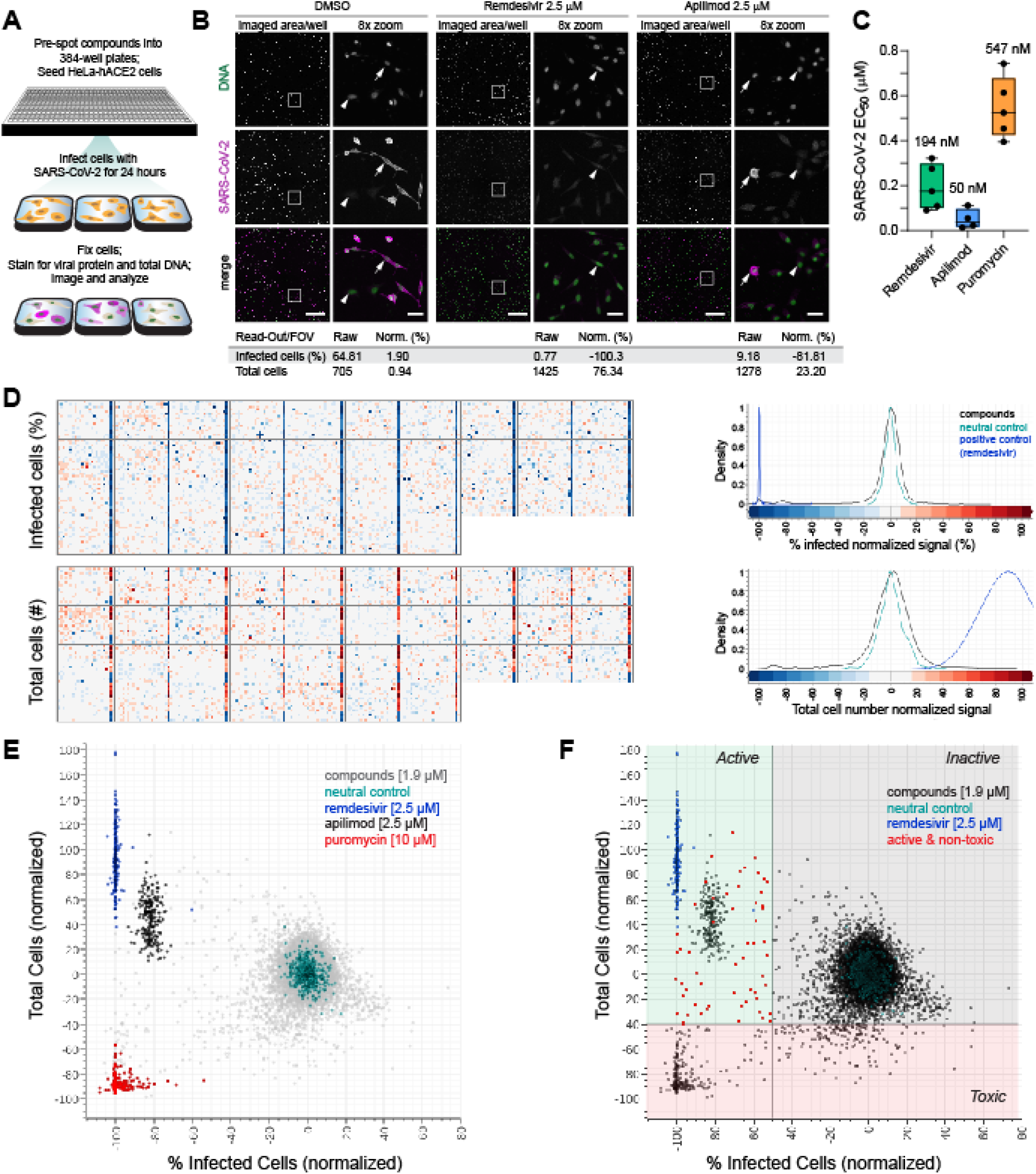
A primary cell-based HCI assay identifies compounds active against SARS-CoV-2 infection. A) Simplified assay workflow. B) Representative images from dimethyl sulfoxide (DMSO)-, remdesivir- or apilimod-treated wells. The entire imaged area per well (4 fields of view taken with a 10× objective and stitched together) is shown for each treatment, as well as an 8-fold magnified segment demarcated with a white box. DNA signal [4’,6-diamidino-2-phenylindole (DAPI)] is colored green, and the virus visualized with immunofluorescence is colored magenta. Infected (arrow) and uninfected (arrowhead) cells are indicated; 500 μm and 50 μm scale bars are shown in the composite and magnified images, respectively. Raw and normalized (Norm.) values calculated from the images is shown. C) Box and whiskers plot of SARS-CoV-2 assay control EC_50_s obtained from independent biological experiments with mean indicated with a bar and all data points shown. Whiskers indicate minimums and maximums. D) Heat map images of normalized data from 1.9 μM ReFRAME screening plates. Normalized activity values for % infected cells and total cell numbers are indicated according to the scale bar and density plot for compound and control wells is shown. DMSO-treated wells are in column 24 and positive control-treated wells (blocks of wells with 2.5 μM remdesivir, 2.5 μM apilimod, or 9.6 μM puromycin) in column 23. Density plots representing the frequency of values associated with each well type are shown on the right. E) Distribution of 1.9 μM ReFRAME screen data for compound and control wells. F) Screen hit selection thresholds.

We validated the assay using compounds with reported activity against Ebola and suspected or previously verified activity against SARS-CoV-2: remdesivir (GS-5734)^10^ (EC_50_ = 194 ± 20 nM; average ± sem of 5 independent experiments) and the PIKfyve inhibitor apilimod (EC_50_ = 50 ± 11 nM, average ± sem of 4 independent experiments) (Fig. 1B). Remdesivir at elevated concentrations was able to eliminate infected cells almost completely (Fig. 1C) and we used it at a concentration of 2.5 μM as a positive control, with data normalized to it and neutral DMSO control wells. While apilimod was more potent than remdesivir, it had a fractionally lower maximal efficacy (85-90% of uninfected cells at the highest effective concentrations) compared to remdesivir. Additionally, we assessed compound toxicity in the context of infection by quantifying the total cell numbers per well, with cytotoxic protein synthesis inhibitor puromycin as a positive control (average EC_50_ = 547 ± 27 nM, average ± sem of 5 independent experiments; HeLa-ACE2 CC_50_ = 2.45 ± 0.23 μM, average ± sem of 5 independent experiments). Notably, a concomitant increase in cell numbers coincided with the antiviral activity of remdesivir and apilimod, likely due to reduction in proliferation of infected cells (Fig. 1B-E). Altering the multiplicity of infection had modest effects on the potency of control compounds in the same experiment, with a 2.7-fold increase in remdesivir’s EC_50_ from MOI=1 to MOI=26, and a 3.7-fold increase in apilimod’s EC_50_, but not that of puromycin (Fig. S1).

Using the developed assay, we ran a pilot screen to assess the activity of 148 small molecules with suspected therapeutic potential against coronavirus infections mined from the available literature^11^ (RZ’ = 0.84). We identified 19 compounds with an EC_50_ < 9.6 μM and, based on data obtained from an uninfected HeLa-ACE2 24-hour live/dead assay, 10 of these were selective (uninfected HeLa-ACE2 CC_50_/SARS-CoV-2 EC_50_ > 10 or uninfected HeLa-ACE2 CC_50_ > 40 μM) (Table S1). This included library/screening lots of apilimod (EC_50_ = 184 nM, CC_50_ > 40 μM) and remdesivir (EC_50_ = 300 nM CC_50_ > 40 μM) that were “rediscovered” in the assay. The higher EC_50_ of apilimod and remdesivir are likely due to slight degradation over time in the screening deck compared to freshly prepared control powder stock.

### Screening ReFRAME, a best-in-class drug repurposing library

Next, we screened the 12,000-compound ReFRAME repurposing library at a final concentration of 1.9 μM and 9.6 μM. Assay quality was maintained throughout both screens (RZ’ of 0.87 and 0.72, respectively) (Table 1) and a clear distinction was apparent in the activity profiles of DMSO vehicle- (neutral control), remdesivir- (positive control), apilimod-, and puromycin- (toxicity control)-treated wells (Fig. 1D, 1E). Hits were selected based on demonstration of >50% reduction in the number of infected cells per well (<-50% activity normalized to neutral controls minus inhibitors) and <40% toxicity based on the total cell number per well (>-40% activity normalized to compound activities, including 10 μM puromycin) (Fig, 1E, 1F) identifying 61 primary hits at 1.9 μM and 266 primary hits at 9.6 μM screening concentrations (hit rates of 0.51 and 2.24%, respectively), with a total of 311 hits.

**Table 1.** Primary and validation screen statistics.

| Library | ReFRAME<br>[1.9 μM] | ReFRAME<br>[9.6 μM] | Total<sup>b</sup> |
| --- | --- | --- | --- |
| Compounds screened | 11,861 | 11,861 | 11,861 |
| Primary hits<sup>a</sup> | 61 | 266 | 311 |
| Average RZ' | 0.8695 | 0.7241 | 0.7968 |
| Hit rate | 0.51% | 2.24% | 2.75% |
| Tested in dose<br>response (DR) | 60 | 265 | 310 |
| EC<sub>50</sub> < 9.6 μM | 43 | 197 | 225 |
| Reconfirmation rate | 71.7% | 74.3% | 72.6% |
| EC<sub>50</sub> < 9.6 μM,<br>CC<sub>50</sub>/EC<sub>50</sub> > 10 | 8 | 41 | 45 |
| Potent and selective<br>hits | 13.3% | 15.5% | 14.5% |

<sup>a</sup> Primary hit thresholds: > 50 % inhibition of infection, < 40% cell toxicity; 6 border-line hits included in 1.9 μM
<sup>b</sup> non-overlapping hits from 1.9 and 9.6 μM screens

The hit rate for the primary screen of the ReFRAME library was high (2.75%), but not unexpected for this collection of bioactive small molecules, many of which are approved drugs or in clinical phases of development and used for a wide assortment of indications (Fig. 2A). To reconfirm and assess potency and selectivity of the primary hits we tested 310 of the available compounds in a 10-point 1:3 dilution dose response format with a top concentration of 9.6 μM. Of these, 225 (72.6%) demonstrated activity with EC_50_ < 9.6 μM against SARS-CoV-2. However, many of the primary screen hits were also cytotoxic, with an unacceptably low selectivity ratio as determined in uninfected HeLa-ACE2 cells (uninfected CC_50_/EC_50_ < 10) (Table 2, Fig. 2B). Because viruses rely on host machinery for replication, it was not unexpected that many of the compounds with antiviral activity also affected vital host processes. Interestingly, this toxicity was sometimes masked in infected cells, as reduction of viral infection by compounds like the protein synthesis inhibitor puromycin and even hydroxychloroquine provided a benefit to cell health in the context of infection but not in uninfected cells (Fig. 2C, Table S1).

**Fig. 2.**
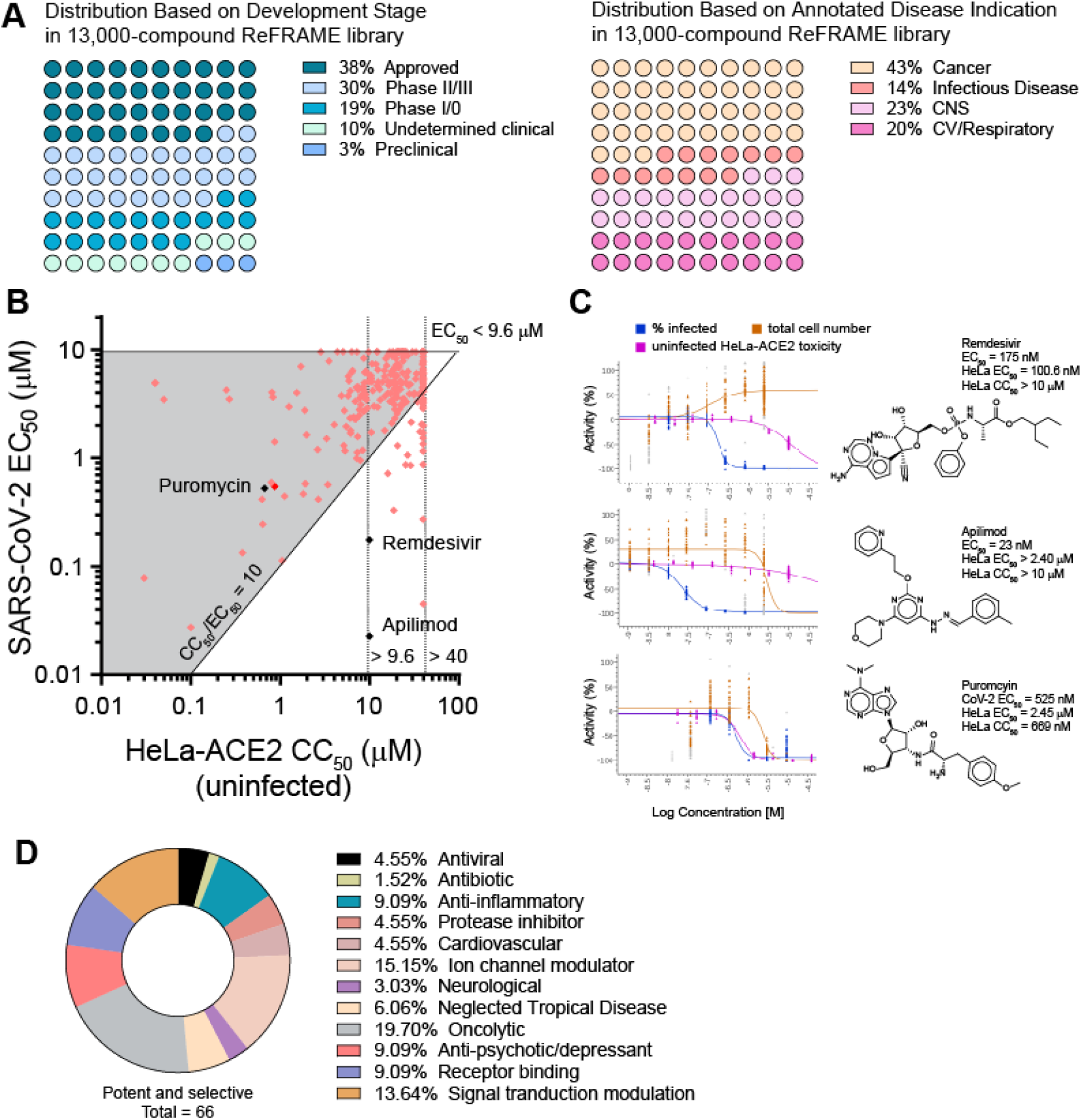
Potent and selective compounds with anti-SARS-CoV-2 activity are identified in the ReFRAME library. A) The composition of the ReFRAME repurposing library with respect to clinical stage of development and disease indication. B) Dose response reconfirmation results, with the SARS-CoV-2 EC_50_ of each compound plotted against its host cell toxicity CC_50_ as assessed in uninfected HeLa-ACE2 cells. Dotted lines represent maximal concentrations tested in dose-response studies for the assay compounds (40 μM) and controls apilimod and remdesivir (10 μM). Activities of controls (black diamonds) and assay compounds (pink diamonds) are shown. Activity of the ReFRAME library copy of puromycin that was screened as part of this hit reconfirmation is also indicated (red diamond). C) SARS-CoV-2 EC_50_ (blue), infected HeLa-ACE2 EC_50_ (orange) and uninfected HeLa-ACE2 CC_50_ dose response curves for the remdesivir, apilimod and puromycin control compounds ran as part of ReFRAME hit reconfirmation. D) Classification of 66 potent and selective compounds according to their functional annotation.

**Table 2.** Attractive reconfirmed hits with activity and selectivity against SARS-CoV-2

| Compound/Drug Name | Target/Mechanism | Clinical Stage | PK Exposure |  | References | SARS CoV-2 EC<sub>50</sub> (μM) | Uninfected HeLa-ACE2 CC<sub>50</sub> (μM) | Synergy score with remdesivir (δ) |
| --- | --- | --- | --- | --- | --- | --- | --- | --- |
|  |  |  | Approx. C<sub>max</sub> | Approx. t<sub>1/2</sub> (hours) |  |  |  |  |
| Remdesivir (GS-5734) | RdRP inhibitor | Emergency FDA registration | 5 μM (human oral) | 1 | 23 | 0.194 | > 10 | *n/a* |
| Hydroxychloroquine | Antirheumatic drug | Registered | 1.7 μM (human oral) | 55 | 24 | 1.30 | 20.43 | -1.572 |
| Halofantrine HCl | Antimalarial; hemozoin inhibitor | Registered | 1 μM (human oral) | 58 | 25,26 | 0.33 | 18.71 | -3.51 |
| Amiodarone | Antiarrhythmic | Registered | 6.8 μM (human oral) | 100 | 27,28 | 1.53 | > 40 | 0.68 |
| Nelfinavir mesylate | HIV protease inhibitor | Registered | 17 μM (human oral) | 7 | 29,30 | 6.22 | > 40 | -1.23 |
| Simeprevir | Hepatitis C NS3/4A protease inhibitor | Registered | 32 μM (human oral) | 16 | 31,32 | 7.82 | > 40 | -0.67 |
| Manidipine | Ca<sup>2+</sup>-channel blocker related to amlodipine | Registered | 10 nM (human, oral) | 3 | 33,34 | 7.97 | > 40 | -4.73 |
| Ozanimod | S1P agonist | Registered | 1.2 μM (human, oral) | 20 | 35 | 4.98 | > 40 | -4.62 |
| Osimertinib | EGFR (Thr790Met Mutant) Inhibitor | Registered | 1.5 μM (human, oral) | 50 | 36,37 | 1.71 | > 40 | *n/d* |
| Diclofensine | Dopamine reuptake inhibitor | Phase III | *unk.*; Generally low for class | *n/d* |  | 6.06 | > 40 | *n/d* |
| Pancopride | 5-HT<sub>3</sub> antagonist | Phase III | 300 nM (human, oral) | 14 | 38 | 4.17 | > 40 | 0.65 |
| Apilimod | PIKfyve inhibitor | Phase II | 600 nM (human, oral) | 6 | 39 | 0.05 | > 10 | 3.57 |
| AQ-13 | Malaria | Phase II | 10 μM (human, oral) | 350 | 40 | 1.07 | 26.6 | -1.77 |
| LY2228820 | p38 inhibitor | Phase II | 5 μM (human, oral) | 190 | 41 | 1.75 | 36.61 | *n/d* |
| MK-2206 | Allosteric AKT inhibitor | Phase II | 571 nM (human, oral) | 80 | 42 | 2.02 | > 40 | 2.09 |
| R-428 | AXL kinase inhibitor | Phase II | 750 nM (oral, humans) | 90 | 43 | 2.88 | > 40 | 7.22 |
| Hanfanchin A | Ca<sup>2+</sup>-channel blocker | Phase I | 0.5 μM (rat, oral) | 21 | 44 | 0.72 | 16.09 | -3.61 |
| L 796568 | Beta-3 adrenergic receptor agonist | Phase I | 0.3 μM (dog, oral) | 13 | 45,46 | 1.15 | > 40 | 3.16 |
| NCO 700 | Cathepsin B inhibitor | Phase I | *unk.* | *unk.* |  | 1.16 | > 40 | 0.21 |
| SMN-C3 | SMA splicing modifier | Phase I | Generally high for class; 2 μM (mouse, oral) | 10 | 47 | 2.00 | > 40 | -0.37 |
| GS-9901 | Phosphatidylinositol 3-kinase delta inhibitor | Phase I | 1.8 μM (rat, oral) | 2 | 48 | 4.13 | > 40 | *n/d* |
| GW-803430 | Melanin-Concentrating Hormone MCH-R1 (SLC-1) Receptor Antagonist | Phase I | 585 nM (rat, mouse oral) | 11 | 49,50 | 4.62 | > 40 | *n/d* |
| SAX-187 | 5 Hydroxytryptamine 6 receptor agonist | Phase I | 0.4 μM (rat, oral) | 3 | 51 | 5.35 | > 40 | -2.98 |
| YM-161514 | Ca-channel blocker | Phase I | *unk.*; Generally low for class | *unk.* |  | 4.53 | > 40 | 0.60 |
| Dutacatib | Cathepsin K inhibitor | Discovery | *unk.* | *unk.* |  | 3.48 | > 40 | 2.95 |
| KC 11404 | Lipoxygenase 5 inhibitor | Discovery | *unk.* | *unk.* |  | 4.41 | > 40 | -2.82 |
| NNC 090026 | Sodium and calcium channel inhibitor | Discovery | *unk.* | *unk.* |  | 7.34 | > 40 | *n/d* |

C<sub>max</sub>, maximum serum concentration; t<sub>1/2</sub>, time to half C<sub>max</sub>; t<sub>max</sub>, time to C<sub>max</sub>; *unk.*, unknown; *n/a*, not applicable; *n/d*, not determined

Between the small pilot and the ReFRAME screen, and not including remdesivir, we identified 67 (66 unique as two different lots of GW-803430 were identified) potent (EC_50_ < 9.6 μM) and selective (CC_50_/EC_50_ >10 or CC_50_ > 39.8 μM) compounds (Fig. 2B, D, Table S1). The top four classes of potent and selective compounds were oncolytic compounds, ion channel modulators, anti-psychotics and receptor binding compounds (Fig. 2D). A fifth of potent and selective hits could be classified as oncolytic drugs, further reflecting the reliance of the virus on host cell processes present in rapidly proliferating cells. The identification of compounds belonging to anti-psychotic, cardiovascular, and even anti-parasitic (neglected tropical diseases) classes may reflect the cationic amphiphilic nature of some of these molecules and their ability to accumulate in and impact acidic intracellular compartments (e.g. late endosomes/lysosomes). Resultant dysregulation of the endo-lysosomal pathway and lipid homeostasis has been suggested to impair viral entry and/or replication^12^ and this mode of action has been speculated for amiodarone and hydroxychloroquine, both identified here as potent and selective hits against SARS-CoV-2 in our screen (Table 2, Table S1). We also identified two selective estrogen receptor modulators (bazedoxifene, EC_50_ = 3.47 μM and raloxifene EC_50_ = 4.13 μM); a class of compounds previously found to inhibit Ebola virus infection^13^.

Out of the identified hits, of highest interest were the newly identified approved oral drugs halofantrine HCl, amiodarone, nelfinavir mesylate, simperevir, manidipine, ozanimod andosimertinib, due to their relatively high exposures or a long history of use as therapeutic agents and therefore potential to be quickly repurposed as COVID-19 treatments following further efficacy vetting in animal models. The viral protease inhibitors nelfinavir and simeprevir have good exposures and based on their described mode of action we speculate that they inhibit SARS-CoV-2 directly. The selective sphingosine-1-phosphate (S1P1) receptor modulator ozanimod is also intriguing as a potential COVID-19 therapy. Selective S1P1 agonists have been shown to provide significant protection against influenza virus infection in murine models by reducing inflammation at the site of infection (reducing release of cytokines by pulmonary endothelial cells and infiltration of lymphocytes into the lungs)^14^, and thus ozanimod may serve as a good combination partner for a direct-acting antiviral drug. The approved calcium-channel blocker manidipine has low exposure but may have the potential to improve COVID-19 disease outcomes for patients. Interestingly, amiodarone has previously been identified as having broad-spectrum antiviral activity in an in vitro screen^15^. Nineteen other compounds in various stages of development such as apilimod (assay control that may inhibit viral entry through disruption of endo-lysosomal trafficking, as found for filoviruses^16^), the protease inhibitors NCO 700 (cathepsin B) and dutacatib (cathepsin K), which may also impact viral entry, all have the potential to show efficacy due to their potency or pharmacokinetic profiles (Table 2). Most of these, except for the very potent apilimod, had modest EC_50_s > 1 μM that did not surpass the potency of remdesivir. However, remdesivir’s requirement for intravenous administration and potentially limited efficacy warrants further investigation into alternative or supplemental therapies. Therefore, we investigated whether the hits identified in our screen would be suitable as partners for a combination therapy with remdesivir.

### Combination therapies

Combination therapies have the potential to increase efficacy of treatment while reducing drug dose of either or both combinations partners, and thus prevent side effects that may be associated with administration of higher doses. Drug combinations can also slow the acquisition of drug resistance. Drug synergy, the increase in activity of the combination therapy beyond what is expected of an additive interaction is rare, yet additive effects themselves have the potential to improve therapy regimens. Conversely, antagonism, the inhibition of activity of the overall combination beyond what would be expected if the compounds acted independently, is an undesirable property. To identify synergistic, additive, and antagonistic interactions between remdesivir and ReFRAME hits, we performed synergy interactions studies in a checkerboard experiment, comparing full dose response of remdesivir against the dose responses of 24 hits with attractive safety and pharmacokinetic profiles in a 10 × 10 matrix (Fig. 2E). We used the *synergyfinder* package^17^ in R to assess the interactions between the tested compounds using the Zero Interaction Potency Model (ZIP)^18^, where a δ score > 10 indicates likely synergy, δ < −10 indicates antagonism, and δ between −10 and 10 suggests an additive interaction. We found no genuine synergy between remdesivir and the compounds tested; however, the combinations were additive, including that of nelfinavir with remdesivir (Fig. 3A, Table 1, Table S1), suggesting these drugs, if they were to prove efficacious in vivo, could potentially be co-administered with remdesivir to increase the overall safety and efficacy of treatment, while limiting the evolution of drug resistance.

**Figure 3.**
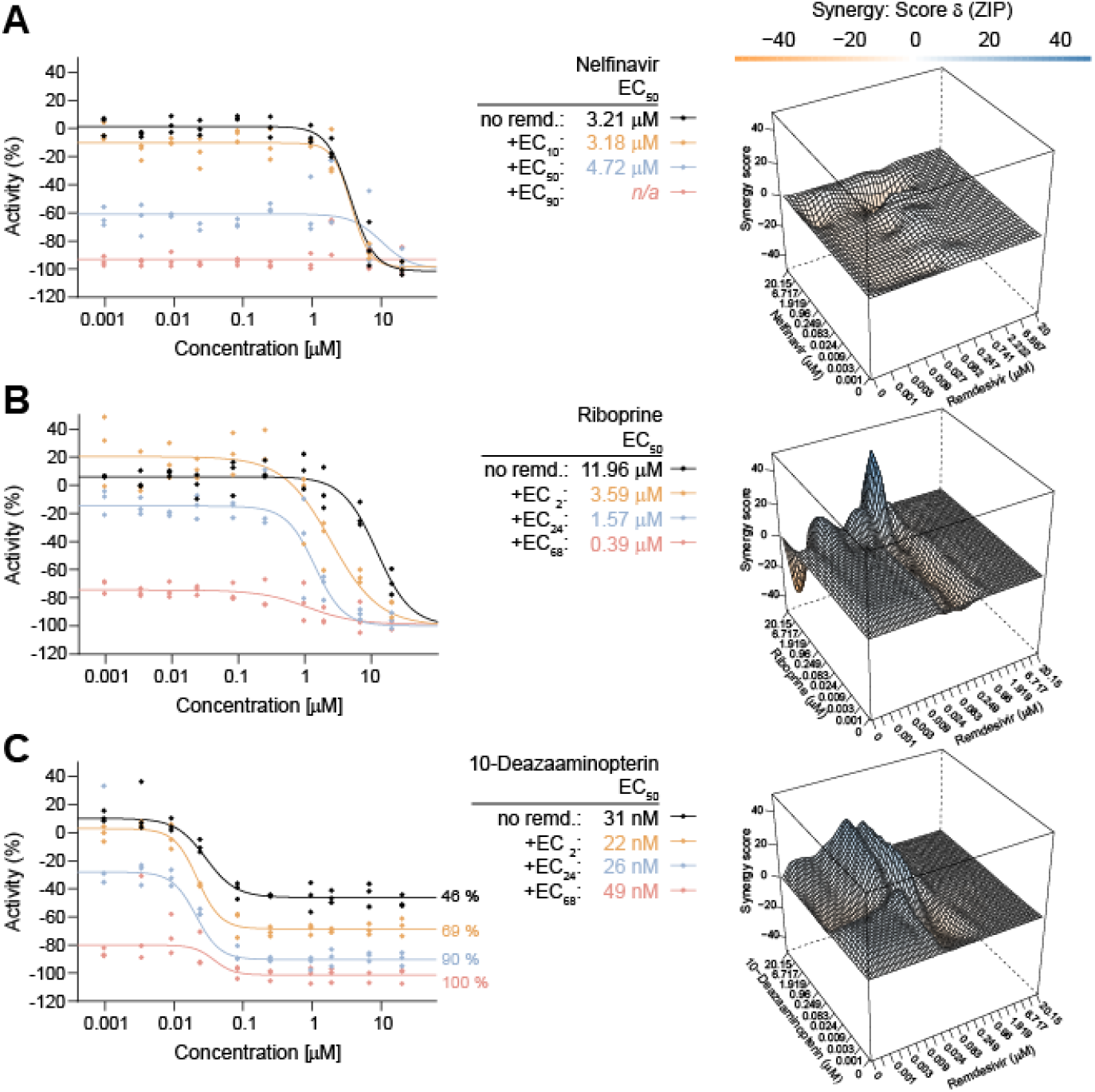
Anti-SARS-CoV-2 activity of drugs in combination with remdesivir showing additive and synergistic interactions. Dose response curves for nelfinavir (A), riboprine (B) and 10-deazaaminopterin (C) without or with increasing concentrations of remdesivir as well as the output of the synergy analysis, a 3-dimensional drug interaction landscape plotting synergy scores across all compound concentrations tested (median scores of 3 technical replicates) are shown. Additive effect: −10< δ <10; synergistic effect: δ >10; antagonistic effect: δ <10. SARS-CoV-2 EC_50_ of each compound in combination with varying concentrations of remdesivir is also shown. Effective concentrations for remdesivir were calculated based on activity of remdesivir in each experiment.

To identify compounds that interact synergistically with remdesivir we carried out a second unbiased ReFRAME screen in the presence of low concentrations (80 nM) of remdesivir. The activity of novel hits from this screen were assessed in the presence and absence of remdesivir. Based on a perceived shift in activity in the presence of remdesivir, compounds were tested in a checkerboard synergy matrix and we identified the nucleoside analog riboprine (N6-isopentenyladenosine, previously investigated as an antineoplastic agent, for treatment of nausea and surgical site infections, and a component of CitraNatal 90 DHA, a prescription prenatal/postnatal multivitamin/mineral tablet) and a folate antagonist 10-deazaaminopterin (an antineoplastic compound currently in Phase II stage of development) as having activities that synergized with those of remdesivir. The synergistic effects for both compounds were observed across specific concentrations, signified as peaks within the 3-dimensional synergy score landscape (Figure 3B, 3C), prompting closer scrutiny of activity of each compound. Riboprine achieved maximal (100%) efficacy over the range of concentrations tested, but addition of EC_2_ of remdesivir shifted its EC_50_ from 12 μM to 3.6 μM, and addition of EC_24_ of remdesivir increased its potency further to EC_50_ = 1.6 μM (Figure 3B). 10-deazaaminopterin showed only 40% maximal efficacy over the range of concentrations tested, but the addition of EC_2_ of remdesivir caused an increase of maximal efficacy from 40% to nearly 65% (where a shift of 2% would be expected) and addition of EC_24_ of remdesivir increased maximal efficacy of the combination from 40% to >80% (Figure 3C). The mechanism of action behind the observed synergies remains to be determined. Riboprine has been reported to block uridine and cytidine import^19^ and through inhibition of protein prenylation to inhibit autophagy^20^ which could impact RNA catabolism^21^ whereas 10-deazaaminopterin has been suggested to inhibit folate dependent enzymes of the purine biosynthesis pathway^19–22^. Therefore, treatment with either riboprine or 10-deazaminopterin may result in reduced intracellular nucleoside pools and in this way synergize with RdRp inhibition by the adenosine nucleoside analog remdesivir. However, another or more direct and specific interaction cannot be excluded and remains to be elucidated. While these findings indicate a promising avenue for further investigation of combination therapies for treatment of COVID-19, the adverse effects of these agents (e.g. inhibition of immunity by 10-deazaaminopterin) would need to be carefully considered in the design and dose selection of in vivo validation experiments.

In summary, by screening the high-value repurposing ReFRAME library we identified 75 unique known drugs or preclinical molecules with activity against SARS-CoV-2 in human cells, 24 of which were tested and showed an additive interaction in combination with the antiviral compound remdesivir recently granted emergency approval for treatment of SARS-CoV-2 infection. We also identified a synergistic interaction between remdesivir and both 10-deazaaminopterin and riboprine. Our data support the advancement of the identified compounds for further profiling in in vivo models to assess their utility (alone or in combination with remdesivir) in combating the COVID-19 pandemic.

## Methods

### Virus generation

Vero E6 cells (ATCC CRL-1586) were plated in a T225 flask with complete DMEM (Corning 15-013-CV) containing 10% FBS, 1×PenStrep (Corning 20-002-CL), 2 mM L-Glutamine (Corning 25-005-CL) overnight at 37 □ 5% CO_2_. The media in the flask was removed and 2 mL of SARS-CoV-2 strain USA-WA1/2020 (BEI Resources NR-52281) in complete DMEM was added to the flask at an MOI of 0.5 and was allowed to incubate for 30 minutes at 34 □ 5% CO_2_. After incubation, 30 mL of complete DMEM was added to the flask. The flask was then placed in a 34 □ incubator at 5% CO_2_ for 5 days. On day 5 post infection the supernatant was harvested and centrifuged at 1,000×g for 5 minutes. The supernatant was filtered through a 0.22 μM filter and stored at −80 □.

### The ReFRAME library: Compound management, drug annotation and screen data access

The ReFRAME library collection consists of nearly 12,000 high-purity compounds (>95%) dissolved in high-quality dimethyl sulfoxide (DMSO). Compound quality control was performed by liquid chromatography-mass spectrometry and/or^1^H-NMR when required. The library was prepared at two concentrations, 2 and 10 mM, to support low-concentration (2–10 μM) and high-concentration (10–50 μM) screening formats. Echo-qualified 384-well low dead volume plus microplates (LP-0200-BC; Labcyte Inc.) were used as the library source plates to support acoustic transfer with an Echo 555 Liquid Handler (Labcyte Inc.). Compounds not soluble in DMSO were plated in water (129 compounds); compounds lacking long-term solubility in DMSO were suspended just before dispensing to avoid precipitation (71 compounds). Additional details available at https://reframedb.org/about.

Associated compound annotation are supported by three widely used commercial drug competitive intelligence databases: Clarivate Integrity, GVK Excelra GoStar, and Citeline Pharmaprojects. As available, annotation data may include status of clinical development and highest stage of development achieved, mechanism of action, drug indication(s), and route of administration.

In accordance with ReFRAME data policies, open access to these data are assured and have been expedited for immediate disclosure at https://reframedb.org/.

### HeLa-ACE2 stable cell line

HeLa-ACE2 cells were generated through transduction of human ACE2 lentivirus. The lentivirus was created by co-transfection of HEK293T cells with pBOB-hACE2 construct and lentiviral packaging plasmids pMDL, pREV, and pVSV-G (Addgene) using Lipofectamine 2000 (Thermo Fisher Scientific, 11668019). Supernatant was collected 48 h after transfection then used to transduce pre-seeded HeLa cells. 12 h after transduction stable cell lines were collected, scaled up and stored. Cells were maintained in DMEM (Gibco, 11965-092) with 10% FBS (Gibco, 10438026) and 1× sodium pyruvate (Gibco, 11360070) at 37□ 5% CO_2_.

### SARS-CoV-2/HeLa-ACE2 high-content screening assay

Compounds were acoustically transferred into 384-well μclear-bottom plates (Greiner, Part. No. 781090-2B). HeLa-ACE2 cells were seeded in 13 μL DMEM with 2% FBS at a density of 1.0×10^3^ cells per well. Plated cells were transported to the BSL3 facility where 13 μL of SARS-CoV-2 diluted in assay media was added per well at a concentration of 2.0×10^6^ PFU/mL (assay multiplicity of infection (MOI) = 2.2). Plates were incubated for 24 h at 34 □ 5% CO_2_, and then fixed with 25 μL of 8% formaldehyde for 1 h at 34□ 5% CO_2_. Plates were washed with 1xPBS 0.05% Tween 20 in between fixation and subsequent primary and secondary antibody staining. Human polyclonal sera diluted 1:500 in Perm/Wash buffer (BD Biosciences 554723) was added to the plate and incubated at RT for 2 h. Six μg/mL of goat anti-human H+L conjugated Alexa 488 (Thermo Fisher Scientific A11013) together with 8 μM of antifade-46-diamidino-2-phenylindole (DAPI; Thermo Fisher Scientific D1306) in SuperBlock T20 (PBS) buffer (Thermo Fisher Scientific 37515) was added to the plate and incubated at RT for 1 h in the dark. Plates were imaged using the ImageXpress Micro Confocal High-Content Imaging System (Molecular Devices) with a 10× objective, with 4 fields imaged per well. Images were analyzed using the Multi-Wavelength Cell Scoring Application Module (MetaXpress), with DAPI staining identifying the host-cell nuclei (the total number of cells in the images) and the SARS-CoV-2 immunofluorescence signal leading to identification of infected cells.

### Uninfected host cell cytotoxicity counter screen

Compounds were acoustically transferred into 1,536-well μclear plates (Greiner Part. No. 789091). HeLa-ACE2 cells were maintained as described for the infection assay and seeded in the assay-ready plates at 400 cells/well in DMEM with 2% FBS and plates were incubated for 24 hours at 37□ 5% CO_2_. To assess cell viability, the Image-iT DEAD green reagent (Thermo Fisher) was used according to manufacturer instructions. Cells were fixed with 4% paraformaldehyde, and counterstained with DAPI. Fixed cells were imaged using the ImageXpress Micro Confocal High-Content Imaging System (Molecular Devices) with a 10× objective, and total live cells per well quantified in the acquired images using the Live Dead Application Module (MetaXpress).

### Data analysis

Image analysis was carried out with MetaXpress (version 6.5.4.532). Primary in vitro screen and the host cell cytotoxicity counter screen data were uploaded to Genedata Screener, Version 16.0.3-Standard. Data were normalized to neutral (DMSO) minus inhibitor controls (2.5 μM remdesivir for antiviral effect and 10 μM puromycin for infected host cell toxicity). For the uninfected host cell cytotoxicity counter screen 40 μM puromycin (Sigma) was used as the positive control. For dose response experiments compounds were tested in technical triplicates on different assay plates and dose curves were fitted with the four parameter Hill Equation. Technical replicate data were analyzed using median condensing. The *synergyfinder* package^17^ in R (version 3.6.3) was used for synergy analysis.

### Data availability

All data are available in the main text or the supplementary materials. Results from the screen of the ReFRAME library have been deposited to the reframedb.org data portal.

## Acknowledgments

We are grateful to Calibr’s Compound Management Group, especially Daniela Tigau, Alonzo Davila, and Hannah Hoang for their assistance with the project, and to Deli Huang at The Scripps Research Institute for supplying the HeLa-ACE2 stably transfected cell line. This work was supported by a grant from the Bill & Melinda Gates Foundation #OPP1107194.

## Author contributions

TFR, CWM, AKC, DRB, PGS, MVH supervised and provided funding for these studies. MAB, CWM and TFR conceived the study and MAB and NB designed and carried out the screening experiments, with MGK, MP, LY, and JR assisting. MAB established image acquisition and analysis. EC and THN handled all compound transfers and provided robotics support. AKG provided chemistry support including quality control on key screening compounds. MVH provided software support. AKC and MVH curated data. MAB formally analyzed and prepared data for publication and MAB, NB, and CWM wrote and edited the manuscript, with contributions from all co-authors.

## List of Supplementary Materials

Fig. S1: Effects of MOI on control compound EC_50_s.

Tables S1: Anti-SARS-CoV-2 activities of known drugs and bioactive molecules identified as potent and selective hits in the pilot and ReFRAME screens.

