## Supplemental Figure 1 for "Oral drug repositioning candidates and synergistic remdesivir combinations for the prophylaxis and treatment of COVID-19"

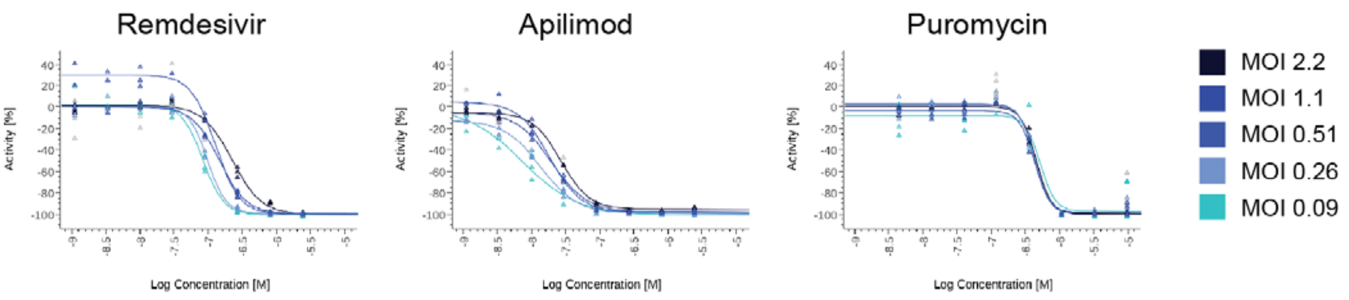


| Remdesivir | Apilimod | Puromycin |
|---|---|---|
| MOI 2.2: 223 nM | MOI 2.2: 27 nM | MOI 2.2: 455 nM |
| MOI 1.1: 145 nM | MOI 1.1: 19.6 nM | MOI 1.1: 431 nM |
| MOI 0.51: 126 nM | MOI 0.51: 18.5 nM | MOI 0.51: 430 nM |
| MOI 0.26: 97 nM | MOI 0.26: 13.6 nM | MOI 0.26: 433 nM |
| MOI 0.09: 83 nM | MOI 0.09: 7.3 nM | MOI 0.09: 520 nM |

**Fig. S1.**

**Effects of MOI on control compound $EC_{50}$s.** Activity of remdesivir, apilimod, and puromycin controls in the SARS-CoV-2/HeLa-ACE2 assay was assessed with MOIs ranging from 0.09 to 2.2. $EC_{50}$ of each compound at the indicated MOI is shown.
